# One-step generation of TCR knock-in mice targeted to the TCRβ locus results in functional mature T lymphocytes

**DOI:** 10.64898/2026.01.13.699338

**Authors:** Jana Bilanovic, Juliana Bortolatto, Susu Duan, Valgerdur Bjørnsdottir, A. Katharina Teetz, Alexandra Thoms, Julian Fishbach, Fabian Schmidt, William Nyberg, Harald Hartweger, Amelia Escolano, Paul G. Thomas, Gabriel D. Victora, Angelina M. Bilate, Johanne T. Jacobsen

## Abstract

Transgenic mouse models expressing predefined T cell receptors (TCRs) have been instrumental in advancing our understanding of T cell biology. However, these traditional models rely on random genomic insertion of large constructs, require labor-intensive embryo manipulation, and frequently result in aberrant TCR expression and phenotypes. These limitations render TCR transgenic models insufficient to meet the mounting demands for rapid and precise model systems to evaluate TCR specificities. To address these challenges, we developed a streamlined method that combines Adeno-Associated Virus (AAV), coupled with CRISPR/Cas9 genome editing to precisely integrate pre-rearranged TCRα/β sequences into the mouse *Trb* locus, enabling the rapid generation of first-of-its-kind TCR knock-in mice with physiological TCR expression and functional T cell differentiation. This approach bypasses the need for technically advanced embryo manipulation and enables rapid generation of models through a universally optimized AAV vector system, significantly enhancing the versatility and utility of monoclonal TCR mice in basic immunology and preclinical research such as cancer immunotherapy and vaccine development, providing a transformative resource to accelerate discovery and translation across disciplines.

## Introduction

Monoclonal T cell receptor (TCR) transgenic mice are indispensable tools in immunological research. They have provided critical insights into the mechanisms of immunological memory and tolerance (Kisielow *et al*, 1988; Sha *et al*, 1988; Teh *et al*, 1988), as well as the pathogenesis of infection (Bot *et al*, 1998; Haanen *et al*, 1999; Kirberg *et al*, 1994; Mitchell & Lawrence, 2003; Oxenius *et al*, 1998; Pircher *et al*, 1990), cancer, and autoimmune diseases (Ashton-Rickardt *et al*, 1994; Bettelli *et al*, 2003; Bruno *et al*, 1995; Cho *et al*, 2020; Haskins *et al*, 1988; Kouskoff *et al*, 1996; Lafaille, 2004; Matsumoto *et al*, 1999; Sebzda *et al*, 1994; Shenderov *et al*, 2021; Verdaguer *et al*, 1997). Despite their fundamental contributions, traditional methods for generating these models are fraught with significant limitations, including inefficiency, variability, and technical complexity, which hinder their widespread application and reliability. To address these challenges, we have developed a technology that enables the rapid, precise, and high-throughput generation of TCR models, paving the way for a more effective interrogation of TCR specificities and their functional relevance.

The first TCR transgenic mice were used primarily to understand central tolerance (Kisielow *et al*., 1988; Sha *et al*., 1988; Teh *et al*., 1988) and thymic selection of CD4^+^ T cells (Berg *et al*, 1989; Kaye *et al*, 1989). This was achieved by inserting pre-rearranged TCRα and TCRβ genes randomly into the genome, using constructs reliant on external enhancers to ensure functional expression (Berg *et al*, 1988; Krimpenfort *et al*, 1988; Teh *et al*., 1988). While subsequent advancements introduced more refined targeting cassettes and regulatory elements (Kouskoff *et al*, 1995; Zhumabekov *et al*, 1995), the inherent randomness of transgene insertion often leads to unintended consequences. These include low expression, often requiring multiple transgene insertions in tandem to obtain sufficient TCR expression (Kouskoff *et al*., 1995). In addition, transgenes can also be deleted, resulting in only endogenous TCR expression in mature T cells (Komori *et al*, 1993). Lastly, the random insertion site of the transgene can lead to aberrant T cell phenotypes, including accelerated onset of T cell lymphomas (Son *et al*, 2018) and other lymphoproliferative diseases (Xiong *et al*, 2013). Because of these stochastic factors, making TCR transgenics requires the generation of multiple founder strains that then must be assayed for proper T cell function. These biological limitations are compounded by the need for pronuclear injection into fertilized oocytes, which requires specialized training and equipment.

The use of CRISPR/Cas9 gene-editing has enabled precise modifications of the TCR loci in somatic cells and significantly advanced T cell-based immunotherapies (Muller *et al*, 2021; Roth *et al*, 2018; Schober *et al*, 2019). An adaptation of such somatic cell orthotopic TCR replacement was utilized for genomic targeting of the *Tra* locus to generate TCR KI mice (Rollins et al, 2023). Genomic *Tra* locus targeting will inevitably allow the selection of mature T cells carrying both exogenous and endogenous Vβ rearrangements due to the nature of TCRα/β assembly mechanisms (Krangel, 2009). Given the dominance of TCRβ in T cell antigen recognition (Boitel et al, 1992; Deckhut et al, 1993; Dillon et al, 1994; Ochi et al, 2015; Zhao et al, 2016), this poses a potential confounding factor as T cells carrying endogenous rearrangements and thus with different antigen specificities can contribute to the functional capacity of such models. Precise modifications of the TCR loci has recently also been accomplished with the generation of a pre-rearranged TCRδ, targeted to the endogenous *Trd* locus (Hahn et al, 2023). This work establishes the physiological expression of a single TCRδ chain paired with a polyclonal repertoire of endogenous chains – an approach well suited for studying repertoire diversity.

To address the challenges of current TCRα/β transgenic approaches, we developed a method that enables the rapid, precise, and high-throughput generation of TCR monoclonal models, allowing for a more effective interrogation of TCR specificities and their functional relevance. Building on CRISPR/Cas9 genome targeting with adeno-associated virus (AAV) delivery we introduced a pre-rearranged CD4^+^ T cell-derived TCRα/β pair into the mouse TCRβ Joining (J)-segment (*Trbj*) locus. The use of an efficient TCR Vβ promoter and removal of endogenous J-segments ensures rapid and physiological expression of TCRs and results in functional monospecificity and efficient allelic exclusion in peripheral CD4^+^ T cells. Founder mice, which are easily identified by PCR, are genetically identical and reliably produce offspring carrying mature monoclonal T cells. These T cells consistently differentiate into T follicular helper (Tfh) or T helper 1 (Th1) subsets in response to antigenic challenge *in vivo*.

Our approach provides a rapid, precise, and cost-effective technique to generate TCR monoclonal mice that meets the need for *in vivo* model systems to evaluate TCR specificities identified from large-scale single-cell analyses or selected for T cell-based cancer therapies (Nowicki *et al*, 2019; Robbins *et al*, 2011; Want *et al*, 2023). This innovation represents a significant advancement in TCR transgenic technology, providing researchers across disciplines with a versatile and efficient tool to explore T cell biology, immune mechanisms, and therapeutic applications and enabling the acceleration of discoveries being translated into clinical solutions for infectious diseases, cancer, and autoimmunity.

## Results

### Generation of TCRα/TCRβ knock-in mice using AAV delivery and CRISPR/Cas9 mediated insertion into the endogenous TCRβ locus

To generate TCR knock-in (TCR KI) mice, we chose Influenza A virus (IAV) as a model infectious agent. To capture a representative selection of IAV-reactive TCRs, we selected a mouse-adapted strain (Webby *et al*, 2003) for which peptide-MHC-II tetramers are available thus facilitating the detection of antigen-specific T cell responses. Wild-type (WT) mice were infected with a mouse adapted strain of IAV, X31, and bronchoalveolar lavage fluid (BALF) was collected 10 days later. We then sorted IAV-specific CD4^+^ T cells from BALF using a PE-labeled MHC-II tetramer consisting of peptide 311–325 from the IAV nucleoprotein (NP_311-325_) in the context of I-Ab (Brincks *et al*, 2013). Single-cell TCRα/β sequencing revealed at least five distinct clonal expansions (Fig. 1A). For gene targeting, we selected the sequences of the most expanded clone which was composed of TRAV6-7/DV9/TRAJ57 for TCRα chain and TRBV3/TRBD2/TRBJ7 for TCRβ chain. We chose this strategy as we aimed to isolate TCR clones that could partake in the local effector response to the infection.

**Figure 1.**
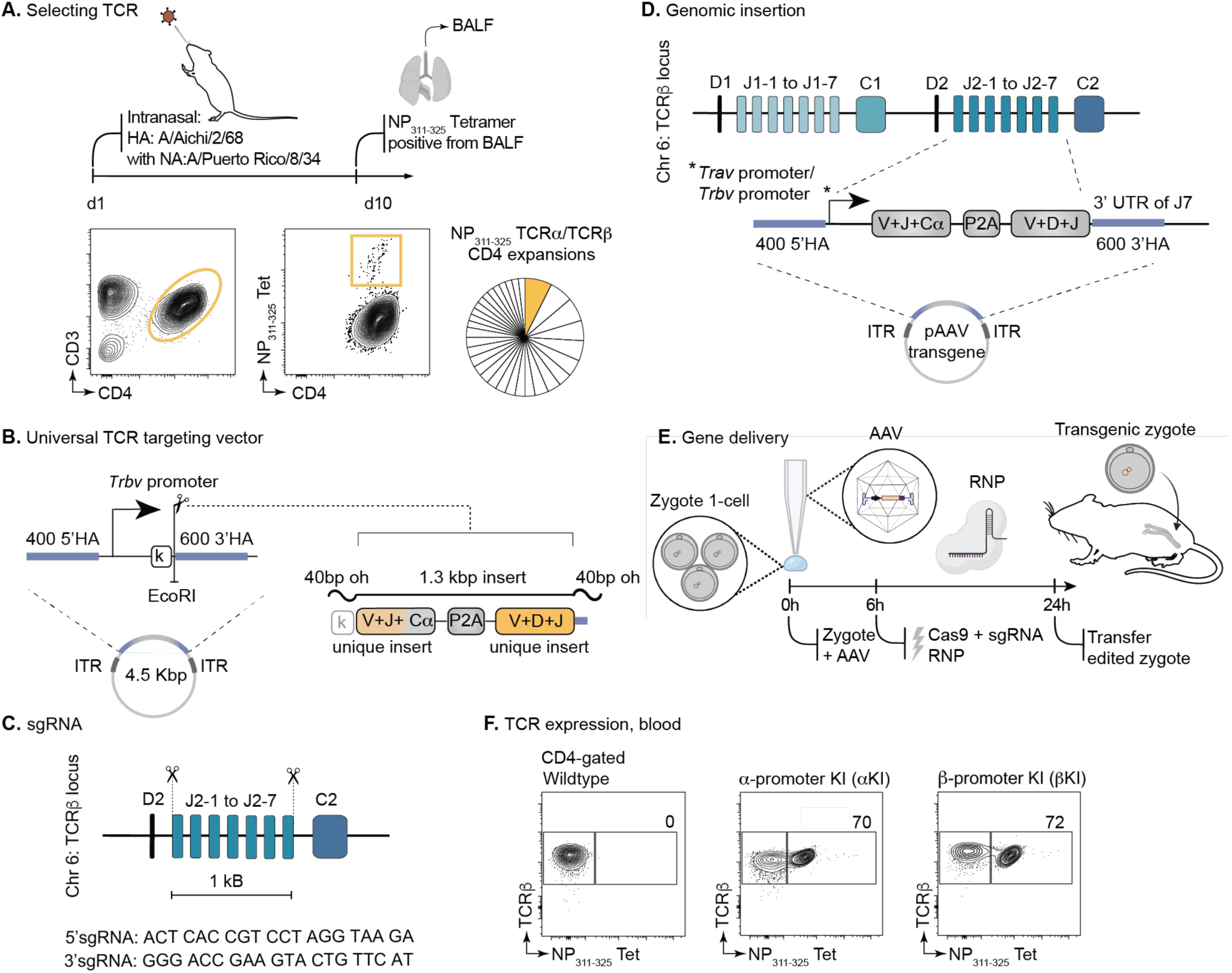
TCR selection and knock-in strategy. A) Selection strategy for the generation of TCR KI mice. Intranasal infection of C57BL/6 with mouse-adapted reverse genetics virus strain (A/Aichi/2/68 HA and NA with A/Puerto Rico/8/34 internal genes), collection of BALF, sorting cells with the influenza nucleoprotein NP_311-325_ tetramer, single-cell processing and selection of expanded TCRα/β sequences. B) Design of a universal targeting vector for TCRα/β expression from the *Trb* locus. AAV vector genomes served as donor templates for homologous recombination into the *Trb* locus by 400 bp homology arms and bicistronic expression under a TCRβ V promoter. Any TCRα/β can be cloned into the universal vector with EcoRI digestion and insertion of ∼1,3 Kbp ds DNA containing the TCR of choice. C) Sequence of sgRNA used to target the *Trb* locus. D) Construct targeting to the *Trb* locus removes J2-1 to J2-7 enabling endogenous J to C2 splicing and bicistronic expression of TCRα/ β under an TCR Vα or Vβ promoter. E) Germline targeting by AAV delivery is facilitated by electroporation 6h post AAV addition to embryos, followed by embryo transfer after 24h. F) Monoclonal TCR expression by blood typing. Targeted mice are screened by NP_311-325_ tetramer staining, and TCRβ antibody. Wildtype mouse is shown as comparison. KI, knock in; HA, homology arms; oh, overhang; AAV, adeno-associated virus; RNP, ribonucleoprotein.

Following our previously reported strategy for generating B cell receptor (BCR) knock-in mice (Jacobsen *et al*, 2018), we inserted tandem TCRα VJ-C and TCRβ VDJ sequences, separated by a P2A ribosome-skipping peptide, into the *Trb* locus. We cloned the tandem TCRα/β sequence into a modified AAV vector (pAAV-MCS), optimized for gene delivery by removing the CMV promoter region and termination region. We further modified the vector with 400-bp homology arms flanking the J-segments of the endogenous mouse *Trbj2* locus and a TCRα TRAV6-7/DV9 promoter upstream of the rearranged TCRs (Fig. 1B). Following zygote transduction and successful genetic recombination, the targeting vector replaced J2-1 – J2-7 in the *Trbj2* locus (Fig. 1C) with a tandem TCRα/TCRβ chimeric gene, which splices onto the endogenous TCRβ C2 upon transcription (Fig. 1D). We also designed the same NP-specific TCR construct using a Vβ promoter directly upstream of the *Trbv* gene segment. We chose the promoter of *Trbv16*, which has clearly distinguishable promoter elements (Anderson *et al*, 1989; Smale & Kadonaga, 2003). Mice constructed with either TCRα or β promoters are referred to as αKI and βKI, respectively.

AAV-mediated gene delivery into the TCRβ locus was performed essentially as previously described (Chen *et al*, 2019) (see methods for details) (Fig. 1E). sgRNAs flanking *Trbj2* were chosen based on high efficiency and low off-target rate using the prediction tool from Integrated DNA Technologies (IDT) (Fig. 1C). Zygotes were incubated with AAVs for 6 h prior to delivery of Cas9 sgRNA ribonucleoproteins (RNPs) by electroporation (Fig. 1E). After an overnight recovery period post-electroporation, healthy zygotes were transferred into pseudopregnant females. The resulting litters were screened by PCR and blood staining to detect NP-specific T cells. To remove potential CRISPR off-target mutations, founder mice were back-crossed for three generations onto C57BL/6J mice prior to experimental use. Both αKI and βKI constructs yielded offspring carrying the KI TCRs in the first electroporation, with 1/4 and 3/18 offspring carrying the Vα and Vβ promoter constructs, respectively. Approximately 70% of circulating CD4^+^ T cells from both KI lines bound to NP_311-325_ tetramer (Fig. 1F), demonstrating efficient integration of the TCR of choice in the *Trb* locus.

### Physiological development of thymocytes and mature T lymphocytes in TCR KI mice

Thymocyte development is a stereotypically ordered sequence of events that results in the production of functional T cells expressing rearranged TCRα and β chains (Ashby & Hogquist, 2024) (Fig. 2A). Conventional TCR transgenics carrying pre-rearranged TCRs display accelerated development of thymocytes into either the CD4 or CD8 single-positive (SP) cells (Kisielow *et al*., 1988). Such accelerated development is also observed in B cells in the bone marrow when B cell receptors are knocked into their respective heavy and light chain loci (Jacobsen *et al*, 2014; Sweet *et al*, 2010). Unsurprisingly, both KI lines showed accelerated thymic development, with most thymocytes in the SP CD4^+^ state and fewer DP thymocytes (Fig. 2B-D). The expression of pre-rearranged TCRα/β likely leads to premature positive selection of thymocytes expressing the NP-specific TCR. Further analysis revealed that a larger fraction of KI CD4^+^ T cells exhibited a mature phenotype (TCRβ^+^CD24^−^) compared to CD4^+^ thymocytes in WT mice (Fig. 2E, F). TCRβ expression, which positively correlates with the maturation of CD4^+^ SP thymocytes, was higher in KI thymocytes compared to WT SP CD4^+^ thymocytes (Fig. 2G).

**Figure 2.**
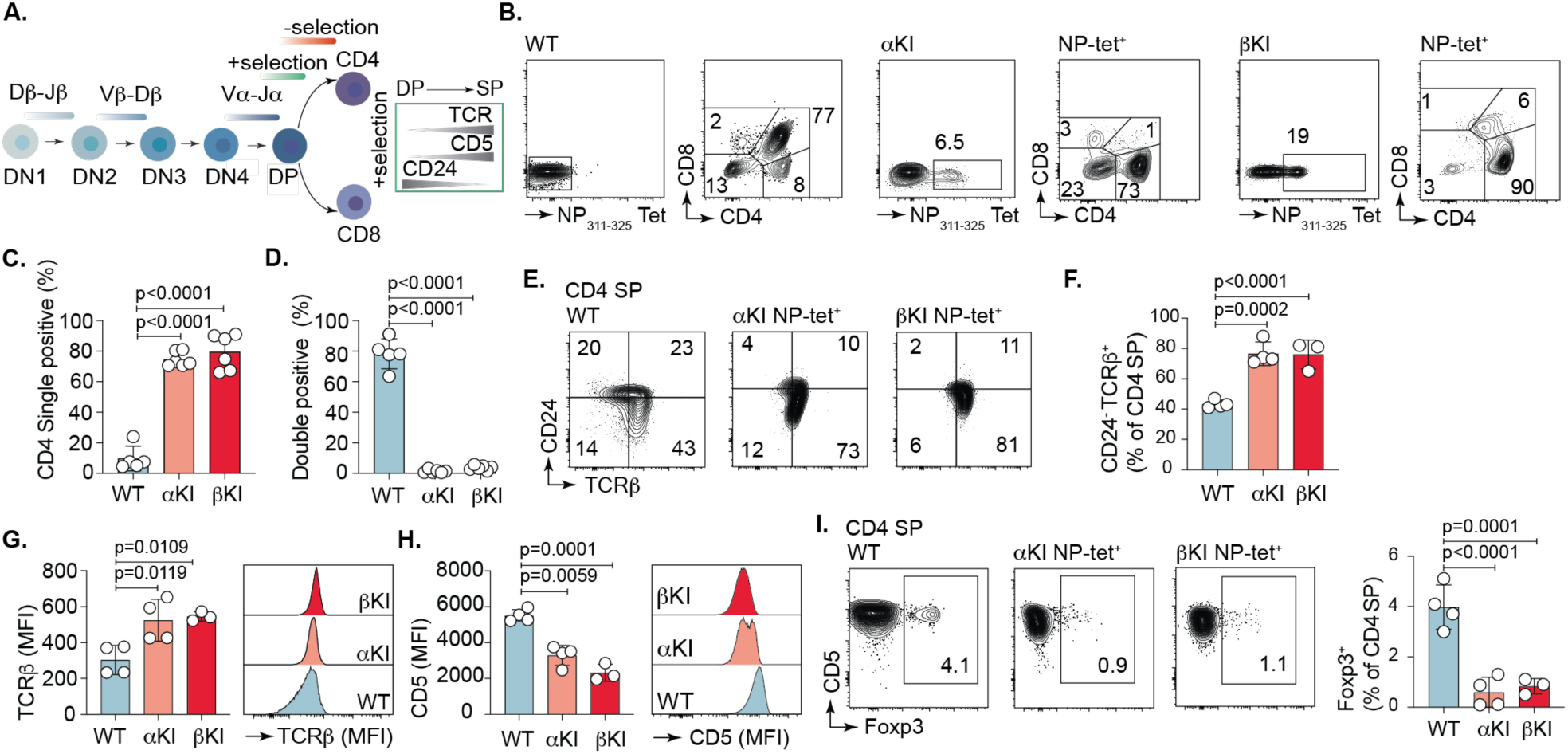
Development of thymocytes in TCR KI mice. A) Schematic depicting thymocyte development progressing through double-negative (DN), double-positive (DP) to single-positive (SP) stages with positive selection accompanied by increasing levels of TCR, CD5 and decreasing level of CD24. B) Subset distribution. Representative flow cytometry analysis of DN, DP, and CD4 or CD8 SP thymocytes for WT, αKI and βKI mice gated on singlets. Subsets are shown for the NP positive fraction of KI mice. C) Quantification of SP state. D) Quantification of DP state. E) Representative flow cytometry plots showing expression of CD24 and TCRβ for CD4 SP thymocytes for WT, αKI and βKI mice. Subsets are shown for the NP positive fraction of KI mice. F) Quantification of mature CD24^neg^/TCRβ^+^ WT CD4^+^ SP thymocytes or KI NP^+^ CD4^+^ SP NP-tetramer^+^ thymocytes. G) Quantification of mean fluorescence intensity (MFI) of TCRβ expression and H) CD5 expression for WT CD4^+^ SP thymocytes or KI NP^+^ CD4^+^ SP thymocytes. I) Representative flow cytometry plots for Foxp3 and CD5 expression in SP CD4 thymocytes. The plots display the percentage of Foxp3^+^ cells within the CD4 SP thymocyte population for WT, αKI and βKI mice. Subsets are shown for the NP positive fraction of TCR KI mice. Graph shows the quantification of Foxp3+ T cells. Data are pooled from four (B-D) or three (E-I) experiments and presented as bar graphs summarizing the results for all experimental groups. All mice were 4 weeks old at the time of analysis. For all plots data and error bars represent the mean ± s.e.m. One way-ANOVA, multiple group comparison (Tukey’s test) was performed.

Expression of CD5 is a read-out for TCR signaling strength, essential for thymic selection (Azzam *et al*, 2001; Pena-Rossi *et al*, 1999) (Fig. 2A). Consistent with the polyclonal nature of the WT population, which includes thymocytes with high-affinity TCRs for self-antigens, we observed higher CD5 expression on WT CD4^+^ thymocytes compared to CD4^+^ KI thymocytes (Fig. 2H). Of note, expression of TCRβ and CD5 was similar between NP-tetramer^+^ and NP-tetramer^−^ CD4^+^ thymocytes among αKI and βKI mice (Suppl. Fig. 1A, B). Self-peptide-MHC interactions with high-affinity thymocytes are known to increase CD5 expression (Azzam *et al*, 1998), dampening TCR signaling and sparing these highly responsive cells from deletion (Azzam *et al*., 2001; Azzam *et al*., 1998; Tarakhovsky *et al*, 1995). In addition, high-affinity self-reactive T cells are selected to become Foxp3-expressing regulatory T cells in the thymus via agonist selection (Hsieh *et al*, 2012). Accordingly, NP-specific CD4^+^ thymocytes showed fewer Foxp3^+^ cells (Fig. 2I). In conclusion, TCR KI mice carry a skewed proportion of mature SP CD4^+^ thymocytes.

### Physiological peripheral development and efficient TCR allelic exclusion for KI T cells

To assess whether TCR KI CD4^+^ T cells can exit the thymus and populate the periphery as efficiently as their WT counterparts, we analyzed spleens from 8-week-old TCR αKI and βKI mice (Fig. 3A). Most CD4^+^ T cells from KI mice exhibited a naïve phenotype, consistent with the absence of the cognate antigen in these mice. Conversely, approximately 12% of WT CD4^+^ T cells displayed a memory/effector phenotype (Fig. 3B, C), a consequence of the polyclonal nature of the WT repertoire and diverse antigen stimulation provided in a specific-pathogen free (SPF) environment.

**Figure 3.**
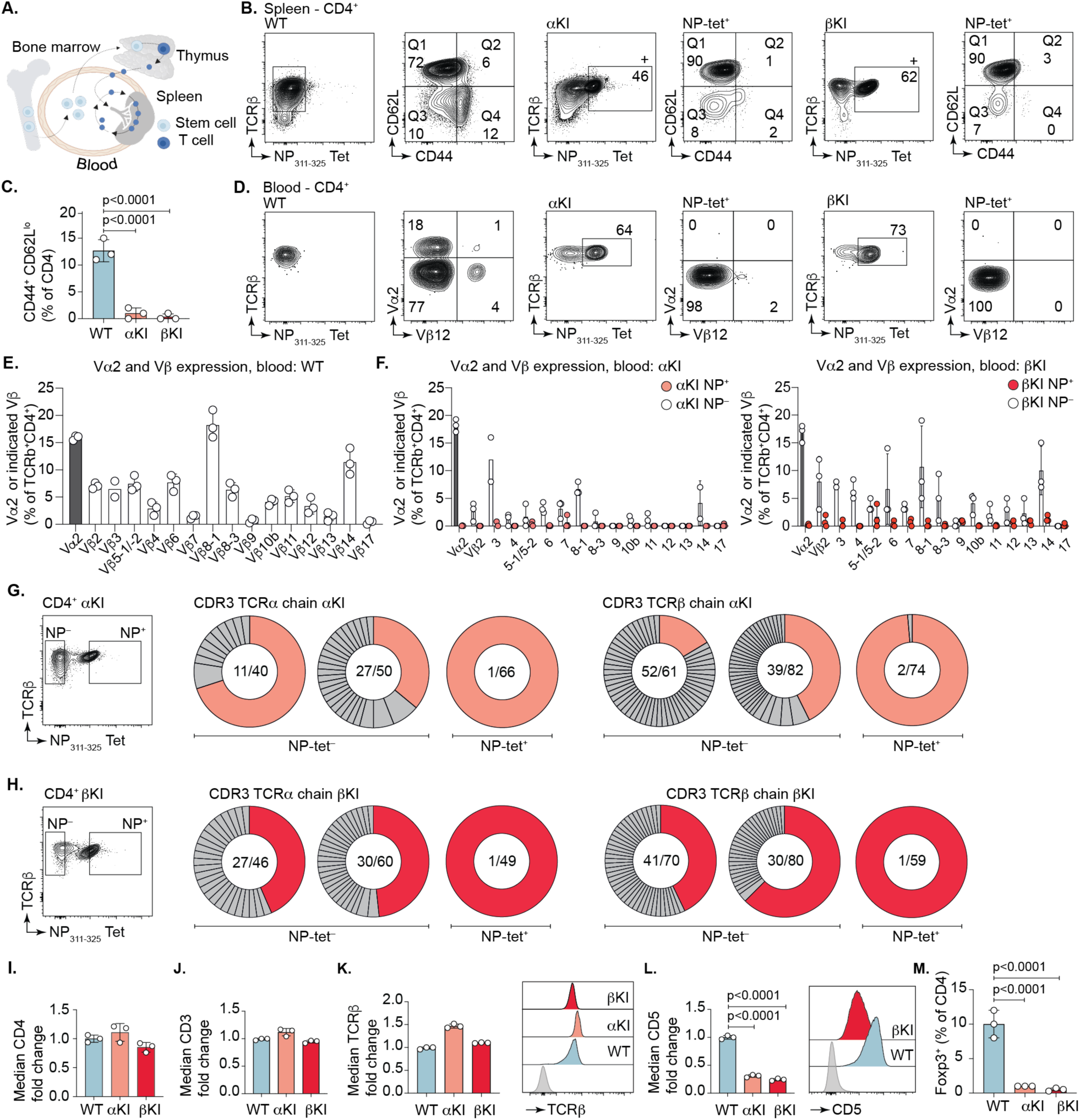
Peripheral development of TCR KI T cells. A) Schematic showing how T cells circulate in blood and populate the secondary lymphoid organs after developing from progenitor stem cells in the bone marrow and passing through thymic development. B) Representative flow cytometry plots showing expression of CD44 and CD62L for CD4^+^ WT, or CD4^+^ NP^+^ KI T cells in spleens of 8 weeks old mice. C) Histogram shows distribution of activated (in gate Q4) T cells. D) Representative flow cytometry plots from WT, NP-KI mice showing TCR Vα2 and TCR Vβ12 expression of CD4^+^ WT or NP^+^ CD4^+^ T cells in blood. E) Expression of TCR Vα2 and TCR Vβ subtypes on a polyclonal CD4^+^ WT repertoire in blood and F) TCR Vα2 and TCR Vβ subtype expression on TCRβ^+^ CD4^+^ NP^+^ or NP^−^ αKI and βKI T cells. Histograms show compiled data from three experiments. Data and error bars represent the mean ± s.e.m. One way-ANOVA, multiple group comparison (Tukey’s test) was performed. G) Single-cell TCRα/β sequencing of splenic T cells from αKI (G) and βKI (H) mice. NP^+^ and NP^−^ CD4^+^ T cells were single cell sorted and TCRα/β rearrangements sequenced. Representative flow cytometry plots show gating strategy. Pie charts indicate unique clones identified/all cells sequenced separated by NP binding. Two mice per group. NP^−^ pie charts are per mouse, NP^+^ pie charts are two mice pooled per group. I-M) Bar graphs show fold change of median fluorescence for CD4^+^ NP^+^ KI T cells over CD4^+^ WT T cells for the indicated costimulatory and activation markers. Histograms indicate MFI of the indicated costimulatory and activation markers for CD4^+^ WT or CD4^+^ NP^+^ KI T cells, or M) percentage of Foxp3^+^ T cells for CD4^+^ WT or CD4^+^ NP^+^ KI T cells. Data are pooled from two (B, C, I-M) or three (D-F) experiments. For all plots unless specifically indicted otherwise data and error bars represent the mean ± s.e.m. One way-ANOVA, multiple group comparison (Tukey’s test) was performed.

We next evaluated allelic exclusion in circulating T lymphocytes by flow cytometry. Silencing of in-frame V-D-J-Cβ genes at the transcriptional and posttranscriptional levels leads to TCRβ allelic exclusion in mouse αβ T cells (Sieh & Chen, 2001; Steinel *et al*, 2010). TCRα allelic exclusion is by default incomplete in TCRα/β T cells, as the TCRα chain can undergo secondary rearrangement on both alleles, resulting in T cells expressing more than one α chain. We stained blood circulating T cells with a panel of 15 FITC-labeled antibodies that target most but not all TCRβ variable families (referred to as “pan-TCRβ”) and with an antibody against TCR Vα2, the most abundant TCRα chain in mice (Fig. 3D). Our KI TCR (*Trbv3*) corresponds to the Vβ16 protein according to standard nomenclature (https://www.imgt.org/IMGTrepertoire/LocusGenes/#J). As there is no antibody available for the Vβ16 TCR, we assessed expression of all other TCRβ families on NP-tetramer^+^ and NP-tetramer^−^CD4^+^ T cells. As expected, the frequency of T cells expressing various TCRβ or TCR Vα2 was overall similar between WT cells and non-NP tetramer-binders (Fig. 3E, F). Conversely, a very low frequency of T cells expressing other TCRβ rearrangements was observed among NP tetramer-binders (Fig. 3F), confirming that both αKI and βKI strains produced NP-specific circulating T cells with the correct TCRα/β rearrangements. Notably, TCR Vα2, which is typically expressed on 15% of all WT CD4^+^ T cells (Fig. 3E), was absent from NP-tetramer binding CD4^+^ T cells from both αKI and βKI mice (Fig. 3D-F), suggesting that most T cells in these two KI strains express the expected rearrangement.

To assess expression of TCRα/β rearrangements at the mRNA level, we performed single-cell TCR sequencing of splenic T cells from αKI and βKI mice and determined the most frequent rearrangement expressed by each cell. As expected, the KI TCRα and TCRβ alleles were dominant in virtually all NP-tetramer binders, indicating efficient allelic exclusion in this population (Fig 3G, H). By contrast, about half of the NP-tetramer non-binders expressed non-KI rearrangements, suggesting that lack of NP-tetramer binding may result from co-expression of one or more endogenously rearranged TCR chains along with the KI pair, possibly leading to mispairing between KI and non-KI chains. Collectively, our results show that the majority of the CD4^+^ T cells in TCR KI mice display the desired monospecific rearrangement, whereas escape from allelic exclusion is readily detectable among NP-tetramer non-binders. Of note, given the low level of escapees with endogenous TCR rearrangement, we observed a small fraction of CD8^+^ T cells and virtually no γδ T cells in the spleens of KI mice (Suppl. Fig. 2A-B).

Successful TCR complex formation and subsequent fine-tuned signaling, requires CD3, the co-receptor CD4, and CD5. Expression of CD4, CD3 and TCRβ by peripheral αKI and βKI T cells were comparable to that of WT cells (Fig. 3I-K). Expression of Nur77—a downstream transcription factor indicative of TCR engagement—was similar between WT and αKI T cells in both Treg and non-Treg cells, suggesting no major impact on TCR signaling (Suppl. Fig. 3A, B). In naïve peripheral T cells, CD5 expression is regulated by TCR contacts with self-pMHC (Kieper *et al*, 2004; Smith *et al*, 2001). Accordingly, we observed that NP-tetramer^+^ KI T cells exhibited reduced CD5 expression (Fig. 3L), mirroring our observations in thymic NP-tetramer^+^ KI T cells and consistent with the low frequency of Foxp3^+^ cells within the CD4^+^ NP-tetramer^+^ T cell population compared to the WT CD4^+^ T cell population (Fig. 3M).

In conclusion, TCR KI mice exhibit normal thymic selection and peripheral CD4^+^ T cell development, with TCR signaling at homeostasis comparable to WT mice. Both KI strains exhibited efficient TCR Vβ allelic exclusion in CD4^+^ NP^+^ T cells. TCRβ allelic exclusion is primarily driven by pre-TCR complex signaling and RAG inhibition (Hoffman *et al*, 1996). Using an endogenous TCRβ promoter likely restricts the window for endogenous recombination more effectively than the TCRα promoter. Our universal targeting vector employs the TRVB16 promoter, emphasizing the importance of promoter choice for optimal TCR expression (see Fig. 1A).

### TCR KI CD4^+^ T cells can differentiate into all major T helper phenotypes

Next, we sought to determine whether KI T cells undergo physiological differentiation into helper T cell subsets upon TCR stimulation (Zhu & Paul, 2010). Such lineage choice is influenced by TCR signal strength (Constant *et al*, 1995; Hosken *et al*, 1995), as well as by costimulatory molecules, the cytokine milieu, and tissue specific cues help determining naïve T cells polarization into helper T cell subsets or states (Chen & Flies, 2013). We first tested the ability of KI T cells to polarize towards different Th subsets *in vitro*. Naïve KI CD4^+^ T cells were stimulated with anti-CD3/CD28 under Th1, Th2, and Th17-polarizing conditions (Sekiya & Yoshimura, 2016) (Suppl. Fig. 4A). CD4^+^ T cells from WT, αKI, and βKI mice, as well as from a previously generated transgenic mouse strain carrying a TCR specific for the same NP peptide (TCR TG) but with a different TCRα/β rearrangement (Jones *et al*, 2023) showed equal capacity to differentiate into T-bet-expressing Th1 cells, GATA-3 expressing Th2 cells, and RORγt-expressing Th17 cells (Suppl. Fig. 4B-D).

To verify that our KI T cells were able to adopt both Th1 and Tfh states efficiently *in vivo*, we used the Influenza-specific TCR TG mouse (described above) as a benchmark due to the well-characterized, robust response of its CD4^+^ T cells to the NP(311-325) epitope in responses to IAV PR8 infection (Jones *et al*., 2023). We first adoptively transferred 5 x10^5^ NP-specific CD4^+^ T cells from NP-TG, TCR αKI or βKI mice into congenically marked recipient mice. Recipients were then immunized subcutaneously in the hind footpad with 5 µg nucleoprotein precipitated in alum, and popliteal lymph nodes were analyzed on day 10 post-immunization (Fig. 4A). Recipients of transferred CD4^+^ T cells from TCR TG and both TCR KI mice developed germinal centers (GCs) and successfully differentiated into Tfh cells (Fig. 4B, C top). Among all Tfh cells, a larger fraction was occupied by transferred T cells (Fig. 4C bottom). Tfh cells generated from TCR TG and TCR KI mice all expressed the lineage defining Tfh transcription factor Bcl6 (Johnston *et al*, 2009; Nurieva *et al*, 2009; Yu *et al*, 2009) to a similar extent (Supp. Fig. 5A). In summary, TCR KI mice can be employed in protein immunization studies.

**Figure 4.**
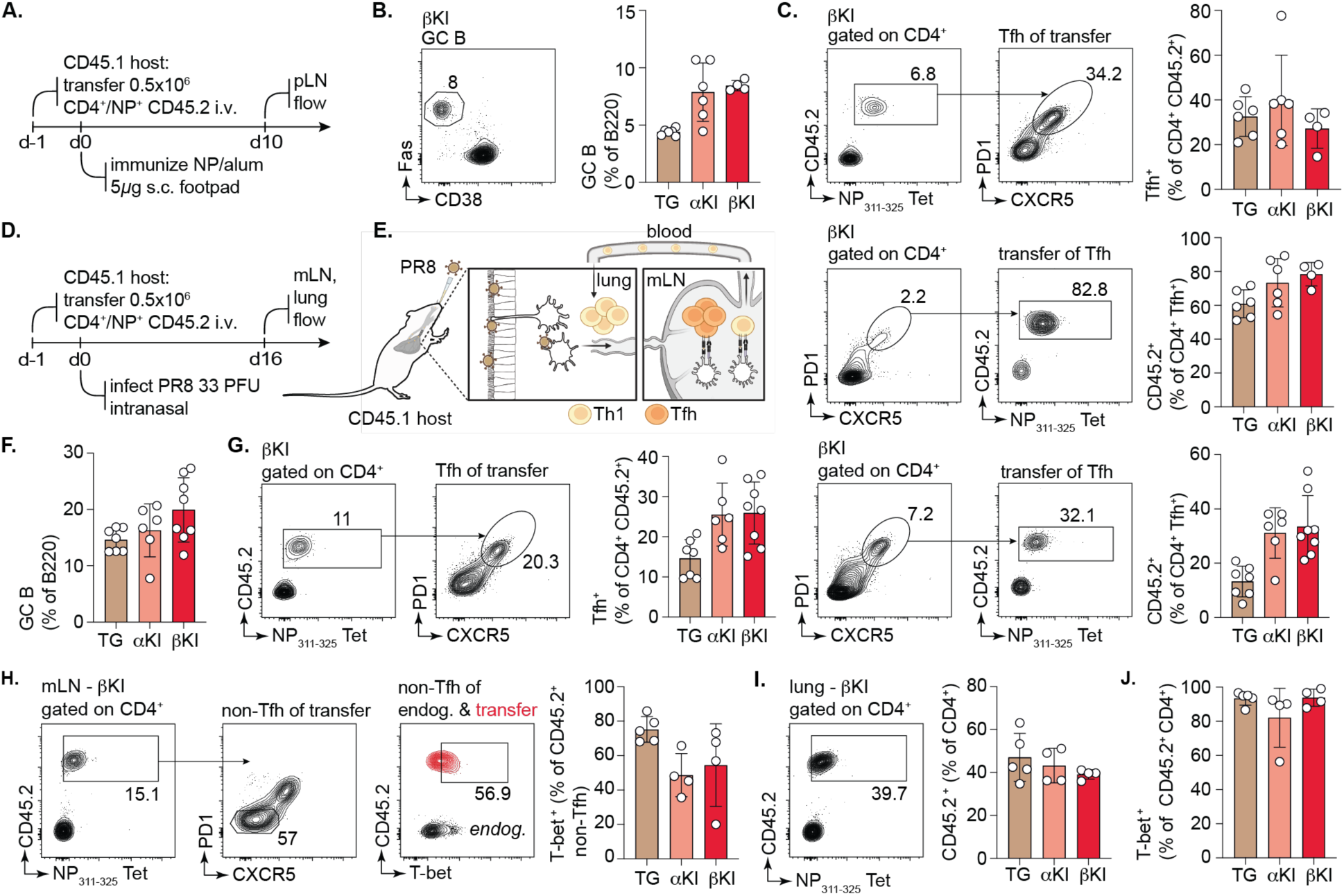
T lymphocytes from TCR KI mice differentiate into Tfh upon protein immunization or Influenza infection. A) Schematic of experimental setup depicting number of NP^+^ CD4^+^ (from CD45.2 donor) and amount of protein used to generate Tfh. pLN; popliteal lymph node. B) Representative flow cytometry plots from a CD4^+^ TCR βKI transfer mouse showing GC B cell gate (gated on B220^+^ B cells) and quantification on the right. C) Representative flow cytometry plots showing Tfh gate out of either all transferred CD4^+^ T cells (top) or transferred CD4^+^ T cells out of the Tfh gate (bottom). Graphs show quantifications. D) Schematic of experimental design for influenza infection experiments. E) Schematic depicting CD4^+^ T cell responses to influenza infection where effector Th1 generated in mediastinal LNs (mLN) will egress through lymphatic vessels, return to blood, and accumulate in lungs by a combination of distinct trafficking signals expressed by airway and alveolar blood vessel endothelial cells. F) Quantification of GC B cells (gated on B220^+^ B cells) in mLNs of infected mice. G) Representative flow cytometry plots on the left show Tfh gates in mLNs of infected mice out of all transferred CD4^+^ T cells. Plots on the right show all transferred CD4^+^ T cells out of the Tfh gate. Graphs show the quantifications. H) Representative flow cytometry plots on the left and middle show the gating strategy for non-Tfh gate (CXCR5^−^/PD1^−^) in mLNs of infected mice. Right plot shows T-bet expression on non-Tfh cells from the host (endog.) or gated on NP^+^ CD45.2^+^ transferred T cells. Graph shows the quantification of T-bet in the transferred cells. I) Representative flow cytometry plots showing NP^+^ CD45.2^+^ transferred T cells in the CD4^+^ T cell compartment of the lungs of host mice. Graph shows the quantification. J) Quantification of T-bet expression in CD4^+^ NP^+^ CD45.2^+^ T cells in lungs of infected host mice. Data are pooled from two (B, C, H, I, J) or three (F, G) experiments. Data and error bars represent the mean ± s.e.m.

We next investigated the ability of NP-specific TCR KI T cells to differentiate into Tfh and Th1 subsets upon PR8 infection. WT mice were transferred with CD4^+^ NP-tetramer^+^ TCR TG or NP-tetramer^+^ TCR KI T cells as above and infected with PR8 the following day. Mediastinal lymph nodes (mLN) and lungs were harvested 16 days post-infection (Fig. 4D, E). Similar to what we observed for protein immunization, mice developed GCs and transferred CD4^+^ T cells efficiently differentiated into Tfh cells in mLNs (Fig. 4F, G). Transferred CD4^+^ KI T cells that differentiated into Tfh cells expressed similar levels of the lineage defining transcription factor Bcl6 (Suppl. Fig. 5B). Both TCR TG and KI T cells expressed T-bet among the non-Tfh population in mLN, indicating they can also differentiate into Th1 cells (Fig. 4H). Furthermore, transferred T cells migrated to and accumulated in the lung tissue where they successfully differentiated into Th1 cells (Fig. 4I-J). Of note, the low numbers of non-tetramer binder escapees present in the TCR KI mice did not expand upon *in vivo* transfer as virtually all recovered KI T cells bound to the NP tetramer (Fig. 4C, G, H, I) and thus do not represent a confounding factor for the outcomes observed. In summary, KI T cells can efficiently differentiate into all major T helper subsets both *in vitro* and *in vivo*. Collectively, these data establish TCR KI mice as a new model to study CD4^+^ T cell responses.

## Discussion

In this study, we establish a rapid and efficient method for the generation of T cell receptor (TCR) knock-in (KI) mice carrying functional monoclonal T lymphocytes. We employed CRISPR-Cas9 and AAV6 gene delivery to precisely integrate pre-rearranged TCRα/β sequences into the endogenous TCRβ (*Trb*) locus, thus ensuring physiological TCR expression. This approach overcomes the limitations of traditional transgenic strategies, such as random genomic integration, associated risks of aberrant phenotypes, and the labor-intensive screening of multiple founder lines, offering a rapid, precise, and cost-effective solution for studying TCR specificity in a physiologically relevant context. Our pipeline leverages universal AAV-based cloning vectors designed for rapid and efficient incorporation of any TCRα/β sequence of interest. Additionally, the electroporation procedure is streamlined to require only basic equipment. This simplicity circumvents the need for extensive technical expertise, improving access to advanced genetic engineering tools. These features enable the rapid and scalable generation of monoclonal TCR models, addressing the growing demand for high-throughput systems to validate findings from single-cell RNA sequencing datasets and to evaluate antigen-specific TCRs in preclinical settings.

A critical determinant of the fidelity of our model is the genomic integration strategy. Linking our pre-recombined TCRβ and TCRα by a 2A peptide enables strict 1:1 stoichiometry of expression, which further enhances the chances of correct assembly of the KI-homodimer, minimizing the risk of orphan KI chains pairing with endogenous chains, and thus enforcing functional monospecificity. While recent approaches have utilized the TCRα (*Tra*) locus for somatic cell targeting (Muller *et al*., 2021; Roth *et al*., 2018; Schober *et al*., 2019) and germline targeting (Rollins et al., 2023), the biological mechanisms governing allelic exclusion differ fundamentally between the *Trb* and *Tra* loci. The *Trb* locus is subject to stringent feedback inhibition, where signaling through a functional pre-TCR complex arrests further Vβ-to-DJβ recombination by allelic exclusion. In contrast, the *Tra* locus lacks this feedback; Vα-to-Jα rearrangement continues on both alleles until positive selection is achieved, creating a permissive window for the generation of dual-receptor T cells. By targeting the *Trb* locus, our model exploits these endogenous developmental checkpoints. We observed that *Trb* KI mice exhibit “accelerated” thymic development, with a skewing of thymocytes toward the mature single-positive (SP) CD4^+^ stage and a reduction in the frequency of double-positive (DP) thymocytes. This enhanced expression of the pre-rearranged TCRα/β complex at the CD4-CD8-double-negative (DN) stage provides a strong positive selection signal that drives rapid transit through the DP compartment. This shortened DP residency time in our KI model would effectively truncate the temporal window available for endogenous *Tra* recombination. This “kinetic exclusion” could lead to reduced endogenous *Tra* rearrangements. This does not exclude that the endogenous TCRα can rearrange in NP^+^ KI T cells. However, we observed that all NP-tetramer^+^ KI T cells carry the functional monospecific TCR. Furthermore, unlike traditional transgenic mice that force allelic exclusion through the non-physiological expression of multi-copy transgenes driven by strong constitutive promoters, our single-copy endogenous promoter strategy exposes the pre-recombined TCR receptor to natural developmental constraints.

Targeting the bicistronic TCR to the endogenous *Trb* locus under the control of a *Trbv* promoter ensures physiological expression by subjecting the transgene to natural developmental checkpoints and transcriptional regulation. Historically, transgenic models have used hybrid strong constitutive promoters to ensure robust, ubiquitous expression of genes (Kim *et al*, 1990; Miyazaki *et al*, 1989). While these promoters address detectability, they introduce biological artefacts arising from overexpression. A critical component of physiological T cell function is the downregulation of surface TCR expression following stimulation, an internalization mechanism that modulates signaling strength and prevents exhaustion. Eyquem et al. (Eyquem *et al*, 2017), showed that constitutive promoters such as the viral LTR or EF1α drive the rapid re-expression of CD19-specific CARs, enforcing prolonged ligand-independent “tonic signaling” and driving cells to acquire an exhausted phenotype. In contrast, placing the receptor under the control of the endogenous *Trac* promoter restored dynamic TCR regulation, enabling T cells to escape exhaustion during chronic antigen stimulation and enhancing anti-tumor efficacy. Similar findings have been reported in murine T cells (Nyberg *et al*, 2023). Thus, driving expression of the TCRα/β complex with an endogenous promoter from the native locus, is essential for generating thymic and peripheral T cells capable of responding physiologically to antigenic stimulation.

The functional competency of the TCR KI mice was validated *in vivo*, with findings highlighting the functional robustness of these TCR KI cells. TCR KI T cells exhibit proper development in both the thymus and periphery, with robust differentiation into T follicular helper (Tfh) cells and Th1 cells across diverse experimental contexts. It did not escape our attention that the ability of TG and TCR KI T cells to differentiate into Tfh or Th1 cells *in vivo* were not identical. These differences could suggest variations in TCR affinity and TCR signaling strength during T cell priming. Strong TCR signals could favor Th1 over Tfh development in TG and KI mice, respectively. However, an impact of TCR targeting to the TCRβ locus on this fate choice cannot be excluded. Both TCR KI and TG cells exhibited Th1 differentiation in the lung, highlighting the dominant role of tissue-specific cues, suggesting that lung microenvironment can override intrinsic TCR properties to dictate lineage commitment. This challenges the dogma that TCR affinity universally dictates T cell fate decisions (Bhattacharyya & Feng, 2020). This observation also underscores how physiological and single TCR expression in monoclonal TCR models is paramount in obtaining accurate data and revealing new biological phenomena. In contrast, less stringent allelic exclusion may result in dual TCRβ expression (Rollins *et al*., 2023), with cells expressing both endogenous and exogenous TCRβ chains. This dual specificity introduces functional heterogeneity and may confound experimental systems where precise antigen recognition is required.

In conclusion, we present a novel, highly efficient and scalable pipeline for generating monoclonal TCR mice, addressing the pressing need for precise and rapid tools to interrogate TCR specificity. Advances in single-cell RNA sequencing have produced extensive datasets identifying TCR sequences linked to immune phenotypes, yet functional validation of these findings requires physiologically relevant and high-throughput models. Likewise, preclinical studies in immunotherapy and infectious disease research increasingly demand scalable systems to evaluate antigen-specific TCRs for therapeutic applications, a gap our approach effectively addresses. By combining precision genome editing with universal cloning vectors and a simplified electroporation protocol, our method enables the rapid generation of multiple monoclonal TCR mouse lines with physiological expression and robust differentiation capacity. This pipeline bridges the gap between large-scale genomic discoveries and functional *in vivo* studies, providing an essential tool for advancing immunological research and accelerating translational applications across oncology, infectious diseases, and autoimmunity.

## Materials and Methods

### Mice

Wild-type C57BL6/J (JAX strain: 000664)*, Nr4a1-EGFP/cre (Nur77-GFP*, JAX strain: 018974*) and* B6.SJL-*Ptprc^a^ Pepc^b^*/BoyJ (JAX strain: 002014) mice were purchased from Jackson Laboratories. C57BL/6J mice (Jackson Laboratory, strain no. 000664). CD1 mice (strain: 482) were purchased from Charles River laboratories. NP Transgenic mice were generously provided by Susan Swain at the UMass Chan Medical School. Knock-in mice were backcrossed to wildtype C57BL6/J mice purchased from Jackson Laboratories for at least three generations. All animal procedures were approved by the Institutional Animal Care and Use Committee of the Rockefeller University (Protocol number 22058-H).

### Isolation of NP-specific T cells from BALF in Influenza infected mice

C57BL/6 mice were anesthetized with avertin i.p. and infected intranasally with the X31 mouse-adapted reverse genetics influenza virus strain (A/Aichi/2/68 HA and NA with A/Puerto Rico/8/34 internal genes). On day 10 following infection, mice were euthanized and a bronchoalveolar lavage was collected, cells isolated, and stained with CD3, CD4, and a tetramer labeled with PE consisting of I-Ab containing the NP_311-325_ peptide.

### Targeting vector generation

AAV serotype 6 has been described previously for mouse generation (Chen *et al*., 2019; Yoon *et al*, 2018). For enhanced delivery a commercial plasmid was modified: pAAV-RC6 Vector (Cell biolabs), was edited with the following custom mutations described previously for their ability to enhance transgene delivery: S663V + T492V (Pandya *et al*, 2015), F129L + Y445F + Y731F (van Lieshout *et al*, 2018) and Y705 + Y731F + T492V (Ling *et al*, 2016). We cloned the tandem TCRα/β sequence into an adeno-associated virus (AAV) transfer plasmid (pAAV-MCS, Biolabs, US catalog number VPK-410), replacing its original promoter and termination region (MluI-PmlI). Following the ITR, we cloned into this backbone 400 bp homology arms immediately 5’of J2-1. The 5’homology arm was followed by a TRAV6-7/DV9 or TRVB16 promoter and the TRAVJ and TRBVDJ separated by a porcine teschovirus-1 2A peptide (Ryan *et al*, 1991) enabling polycistronic and stoichiometric expression of TCRα and TCRβ recombined genes. Directly following the J7 is the endogenous J2-7 3’UTR which also served as our 600 bp 3’ homology arm, flanked 3’ by the ITR of the AAV6 backbone. We constructed a universal TCRβ targeting construct, where any pre-recombined TCRα/TCRβ can be inserted by Gibson assembly into the single site EcoRI digested backbone. Mice were genotyped with a forward primer binding in the Vα gene and a reverse primer binding in the P2A sequence (5’ACT CAC CGT CCT AGG TAA GA 3’ and 5’GGG ACC GAA GTA CTG TTC AT 3’respectively), resulting in a 570 bp fragment upon positive integration.

### Production and Purification of Recombinant AAV6 Particles

HEK-293T cells (ATCC, cat. no. CRL-3216) were cultured in high-glucose DMEM (Thermo Scientific, cat. no. 11965092) containing 10% fetal bovine serum (Thermo Scientific, cat. no. A3382101), 1 mM sodium pyruvate (Thermo Fisher, cat. no. 11360070), 1% Antibiotic-Antimycotic (Thermo Fisher, cat. no. 15240062), and cultured at 37 °C in a humidified atmosphere containing 5% CO₂. For the production of adeno-associated virus (AAV), AAV293 cells were plated in15 cm tissue culture dishes and grown to approximately 90–95% confluence.

Recombinant AAV particles were produced using a triple plasmid transfection, using the pHelper plasmid (Cell Biolabs; 11,854 bp), a modified pAAV-RC6 plasmid encoding AAV6 Rep and Cap genes (7265 bp), and a pAAV transfer plasmid containing a tandem TCRα/β expression cassette (5842 bp). Stock solutions of the plasmids were prepared in TE buffer at 1 mg/mL and combined in a 1:1:1 molar ratio, resulting in a total DNA mass of 91.4 μg per 15 cm dish. Polyethylenimine (PEI, 1 mg/mL; Fisher Scientific, cat. no. 24765-100) was used for transfection at a ratio of 4 μL PEI per 748 μg of DNA. The PEI and plasmid DNA were diluted separately in serum-free DMEM, incubated at room temperature for 10 minutes, mixed together, and incubated for another 20–30 minutes to facilitate complex formation. This mixture was then added dropwise to the cell cultures. To reduce cytotoxicity caused by PEI, the growth medium was exchanged for DMEM supplemented with 2% FBS after 14–16 hours. The cultures were maintained for three days following transfection. On the third day, cells were harvested using 0.05% Trypsin-EDTA (Thermo Scientific, cat. no. 25300054), and both the cells and culture supernatant were collected. Cellular debris was removed by centrifuging the combined material at 2000 g for 10 minutes at 4°C. The supernatant was clarified by filtration through a 0.2 μm PES membrane (Thermo Scientific).

Viral precipitation was achieved by adding a 5× PEG/NaCl solution (400 g PEG 8000 and 24 g NaCl dissolved in 1 L Milli-Q water, pH 7.4, autoclaved) to the supernatant at a 1:4 volume ratio. This mixture was incubated overnight at 4 °C. The viral particles were then pelleted by centrifugation at 2500 g for 1 hour at 4 °C and resuspended in 1 M HEPES buffer (Thermo Scientific, cat. no. 15630080). An equal volume of chloroform was added to the resuspended pellet after thorough vortexing. The sample was vortexed vigorously for 2 minutes and then centrifuged at 1000 g for 5 minutes at room temperature. The aqueous phase, which contained the AAV particles, was carefully extracted to a new polypropylene tube without disturbing the chloroform layer. Any remaining chloroform was allowed to evaporate under a chemical hood for 30 minutes. The AAV solution was sterilized using a 0.22 μm filter (Millipore Sigma, cat. no. SLGPM33RS) before undergoing buffer exchange and concentration with 100K MWCO Amicon Ultra centrifugal filters (Millipore Sigma, cat. no. UFC801024). To prevent viral loss through membrane adsorption, filters were pre-treated with 10% Pluronic acid (Thermo Scientific, cat. no. 24040032) for one hour and equilibrated with formulation buffer (DPBS with 0.001% Pluronic acid). The volume was reduced by ≥200-fold through iterative centrifugation at 3200–3500 g at 15 °C until the target concentration was reached. Residual plasmid DNA was degraded by treating the final preparation with DNase I (New England Biolabs, cat. no. M0303L). Viral titers were determined via SYBR Green real-time qPCR (PowerUp SYBR Green Master Mix, Thermo Fisher, cat. no. A25741) using primers targeting AAV inverted terminal repeats (ITR) (forward: 5’-GGAACCCCTAGTGATGGAGTT; reverse: 5’-CGGCCTCAGTGAGCGA) and a titrated AAV6 standard (Charles River RS-AAV6-FL)^24^. The final AAV product was aliquoted (10–20 μL) and stored at −80 °C.

### Zygote Manipulation

To generate fertilized zygotes for a single electroporation session, 20 female C57BL/6J mice (5–6 weeks old) were superovulated via intraperitoneal injection of 5 IU Pregnant Mare Serum Gonadotropin (PMSG, ProSpec, HOR-272), followed by 5 IU human Chorionic Gonadotropin (hCG, Sigma-Aldrich, C1063-10VL) 48 hours later. These females were mated overnight with 20 C57BL/6J males (8–10 weeks old). The following morning, mice were checked for copulation plugs, and zygotes were harvested approximately 12 hours post-coitum. For that, females were euthanized by cervical dislocation, and oviducts were dissected to release zygotes into two 100 μl drops of pre-warmed EmbryoMax M2 Medium containing Phenol Red and Hyaluronidase (Sigma-Aldrich, cat. no. MR-051-F) in a non-treated petri dish. Embryos were incubated in this medium until cumulus cells detached and the embryos separated from aggregates. Using a mouth-controlled fine glass pipette, denuded zygotes were washed in four sequential 30 μl drops of fresh, pre-warmed M2 medium without Hyaluronidase (Sigma-Aldrich, cat. no. MR-015-D). The embryos were then transferred into pre-warmed 30 μl droplets of EmbryoMax KSOM (Sigma-Aldrich, cat. no. MR-106-D) containing 2x10⁹ AAV genomic units, overlaid with mineral oil (Sigma-Aldrich, cat. no. M8410-100ML). These KSOM droplets containing AAV had been pre-incubated for at least 30 minutes at 37 °C in 5% CO₂ and 95% humidity. Groups of 40–50 embryos were added to each droplet and incubated at 37 °C for 5–6 hours.

Following incubation and prior to Cas9/sgRNA RNP electroporation, zygotes were washed three times in Opti-MEM droplets (Fisher Scientific, cat. no. 51985034) using a mouth-pipette. Subsequently, 20–25 embryos were moved to 10 μl of electroporation solution mixed with Cas9/sgRNA RNPs, placed between the electrodes of a BEX electroporation chamber (Bex, cat. no. CUY21EDIT2). The electroporation parameters were set as follows: 25 V pulse voltage (“PdV”), 300 mA amplitude (“pDA”), 3 ms pulse duration (“Pd on”), 97 ms interval (“Pd off”), and 3 pulses (“PdN”) with logarithmic current decay. After electroporation, the embryos were returned to the mineral oil-covered KSOM droplets containing AAV and cultured overnight at 37°C with 5% CO₂^38^. The following day, 2-cell stage embryos were washed 10 times to clear residual virus and selected for transfer^39^. Approximately 30 embryos were transferred into pseudopregnant CD1 females^40^. The surrogate mothers were CD1-elite albino females (6 weeks old), which had been mated the previous night with vasectomized CD1-elite males (8 weeks old at the time of surgery, with a 3-week recovery period) to induce pseudopregnancy. Roughly 30 embryos were implanted into the oviduct of each surrogate female.

### Cas9/sgRNA preparation

Alt-R™ S.p. Cas9 Nuclease V3 and sgRNA were obtained from (Integrated DNA Technologies, IDT). sgRNA was resuspended in nuclease-free Duplex buffer (IDT, cat. no. 11-01-03-01) to 100 µM and stored in single use aliquots of 5 μl in -80. Cas9 is provided in solution at 10 µg/µl. Prior to electroporation Cas9/sgRNA RNPs was prepared using a total of 1.2 μl of sgRNA (from the 100 μM pre-aliquots), mixed with 1.3 μl Cas9 together with 7.5 μl of electroporation buffer (20 mM HEPES, pH 7.5, 150 mM KCl, 1 mM MgCl2 and 10% glycerol in water).This was incubated for 10 minutes at 37°C before diluting 1:1 with Opti-MEM, which is the solution in which embryos are electroporated.

### Adoptive cell transfers

Spleens were harvested and homogenized through at 70 µm cell strainer. Red blood cells were lysed using ACK buffer (Fisher Scientific, cat. no. A10492-01). CD4^+^ T cells were purified by negative magnetic-activated cell sorting (MACS) using a mix of biotinylated antibodies targeting Ter119, CD11c, CD11b, CD25, B220, NK1.1 and CD8, followed by anti-biotin beads (Miltenyi Biotec, cat. no. 130-090-485), following manufacturer’s instructions. After counting, cells were normalized to NP-binders and 5x10^5^ NP^+^ CD4^+^ T cells were transferred intravenously in 100 ul of sterile PBS 1X (Phosphate-buffered saline, Gibco, cat. no. 10010023). Cells were normalized to NP-binders based on percentage of NP^+^ CD4^+^ T cells in peripheral blood of mice using NP_311-325_ tetramer staining (provided by the NIH Tetramer core). Each NP mouse used was blood-typed before cell harvest to normalize NP binders.

### Immunization and infection

To induce GC formation in popliteal lymph nodes, mice were immunized with subcutaneous injections into the hind footpad with 5 ug of recombinant NP protein (Sino Biological, cat. no. 11675-V08B) precipitated in alum (Imject Alum, Thermo Scientific, cat. no. 77161) at a 1:1 antigen in PBS:alum ratio (v/v) with a final volume of 25 ul. For influenza infection, mice were anesthetized with a ketamine/xylazine mix diluted in sterile PBS 1x and infected intranasally with 33 PFU of mouse-adapted influenza PR8 virus (provided by M. Carroll, Harvard Medical School).

### In vitro Th differentiation assay

T cells were isolated as described in “Adoptive cell transfers”. Naïve CD4^+^ T cells were differentiated into Th1, Th2 and Th17 subsets following the “T helper cells: a complete workflow for cell preparation, isolation, stimulation, polarization and analysis” protocol from Miltenyi Biotec with small modifications. In short, 2.5x10^5^ naïve CD4^+^ T cells were cultured with CD3/CD28 Mouse T-Activator Dynabeads (Gibco, cat. no. 11456D) at a 1:3 ratio in T cell media in 96 well plates. For Th1 differentiation, cultures were supplemented with 10 ng/ml IL-12, 10 ng/ml IL-2, 10 ug/ml Anti-IL-4 antibody. For Th2 differentiation, cultures were supplemented with 10 ng/ml IL-4, 10 ng/ml IL-2, 10 ug/ml anti-IFN-γ antibody. For Th17 differentiation, cultures were supplemented with 20 ng/ml IL-6, 10 ng/ml IL-23, 10 ng/ml IL-1β, 2 ng/ml TGF-β, 10 ug/ml Anti-IL-4 antibody, 10 ug/ml anti-IFN-γ antibody. For Th0 differentiation, cultures were supplemented with 10 ng/ml IL-2. On day 2 of the culture, cells were pipetted up and down to break up cell aggregates. On day 3, cells were supplemented with fresh T cell media at a 1:1 ratio. For Th1 and Th2, the media was supplemented with 10 ng/ml IL-2. On day 4, cells were split 1:2 and Th1 and Th2 were supplemented with additional 10 ng/ml IL-2. For Treg differentiation, 2x10^6^ naïve CD4^+^ T cells were plated in 6-well plates and supplemented with CD3/CD28 Mouse T-Activator Dynabeads (Gibco, cat. no. 11456D) at a 1:1 ratio, 200 U/ml IL-2 and 4 ng/ml TGF-β. On day 5, T cell cultures were harvested for FACS analysis of intracellular transcription factors according to manufacturer’s instructions.

### Isolation of lymphocytes

Circulating lymphocytes were collected via submandibular blood collection using sterile lancets. Blood was collected in microfuge tubes with 20 ul 1M EDTA (Invitrogen, cat. no. 15575020). Spleens were harvested and homogenized through at 70 µm cell strainer. Red blood cells from blood and spleen were lysed using ACK buffer (Fisher Scientific, cat. no. A1049201) prior to staining for flow cytometry. Popliteal and mediastinal lymph nodes as well as thymus were collected in microfuge tubes with 100 ul of sterile PBS 1X and mechanically disassociated into cell suspensions using disposable micropestles (Avanot, cat. no. AXYGPES-15-B-SI). Cell suspensions were filtered through 70 µm cell strainer before further handling. Lungs were collected into 12-well plates with 1.5ml of HBSS (Gibco, cat. no. 24-020-117) supplemented with CaCl_2_, MgCl_2_, 1.5 mg/ml Collagenase D (Roche, cat. no. 11088858001) and 1 mg/ml DNase I (Roche, cat. no. 10104159001) and dissociated into small pieces using scissors. Lungs were digested with the enzyme mix for 45 min at 37 °C and 80 rpm shaking in a shaker. Next, the suspension was pipetted up and down around 15 times in a 5 ml stripette to break down clusters and then filtered through a 70 µm cell strainer. Immune cells were isolated using a Percoll gradient (Cytiva, cat. no. 17-0891-01). Red blood cells and debris were discarded and the interface containing lymphocytes washed with 1X PBS and filtered again through a 70 µm cell strainer before further handling.

### Flow cytometry

Isolated lymphocytes were resuspended in 1X PBE (PBS 1X supplemented with 0.5% BSA (Sigma-Aldrich cat. no. A9647-500G), 1mM EDTA) and first incubated in 1 µg/ml of anti-CD16/32 (BioXCell, cat. No. BE0307) for 5 min at room temperature. After washing with PBE, cells were stained with surface antibodies diluted in PBS 1X for 40 min on ice. For intracellular staining of transcription factors, cells were handled following manufacturer’s instructions using the Foxp3/Transcription Factor kit (eBioscience, cat. no. 00-5523-00). Cells were washed with 1X PBE and filtered through a 70 µm filter before acquisition. Samples were acquired on a FACSymphony A5 (BD Bioscience). Data was analyzed using FlowJo v10.10.

### Single-cell TCR sequencing

Single cells were index-sorted using a FACS Symphony S6 into 96-well plates containing 5μL of lysis buffer (TCL buffer, QIAGEN 1031576) supplemented with 1 % β-mercaptoethanol and frozen at −80 °C prior to RT-PCR. RNA and RT-PCRs for TCRα and TCRβ were prepared as previously described (Bilate *et al*, 2020; Dash *et al*, 2011; Han *et al*, 2014). PCR products for TCRα and TCRβ were multiplexed with barcodes and submitted for MiSeq sequencing using True Seq Nano kit (Illumina) (Han *et al*., 2014). Fastq files were de-multiplexed and paired-end reads were assembled at their overlapping region using the PANDASEQ (Masella *et al*, 2012) and FASTAX toolkit. Demultiplexed and collapsed reads were assigned to the wells according to the barcodes. Fasta files from MiSeq sequences were then aligned and analyzed on IMGT (http://imgt.org/HighV-QUEST) (Brochet *et al*, 2008). In-frame TCRα and TCRβ junction sequences were included in the analysis. Cells sharing identical CDR3 aminoacid sequences were considered the same clone. For each cell, CDR3α/β sequences were ranked by read abundance, and cells were retained only if the dominant sequence exceeded a minimum read threshold (>10 reads), showed internal redundancy, differed from secondary sequences by at most a single terminal aminoacid substitution, or met a dataset-wide abundance criterion. Filtered TCRα and TCRβ CDR3 sequences were collapsed into unique clones and quantified per sample to generate pie charts.

### Statistical analysis

Statistical tests were carried out in GraphPad Prism 10.0 software and edited for appearance using Adobe Illustrator 27.1.1. Multivariate data were analyzed by one-way analysis of variance with Tukey’s post-hoc tests for pairwise comparisons.

## Acknowledgements

We thank the members of the Victora Lab for helpful discussions and advice; D. Mucida for critical review of the manuscript; M. Nussenzweig for input in the use of AAV for transgenesis; the Comparative Bioscience Center for mouse housing and care; S. Swain at the University of Massachusetts for sharing the transgenic NP TCR mouse with us; the NIH Tetramer Core for providing us with the NP311-325 tetramer; M. Carroll at Harvard Medical School for providing us with the mouse-adapted influenza PR8 virus; S. Yin for writing a script to analyze the single-cell TCR sequencing data; and all Rockefeller University staff for their continued support. This study was supported by a European Research Council Starting grant 101042650 to J.T.J.. G.D.V. is an HHMI investigator. J.B. was supported by a Boehringer Ingelheim Fonds PhD fellowship. A.K.T. was supported by a scholarship from the German National Academic Foundation. H.H. was supported by a Hemophilia Association of New York research grant. Some schematics were created using Biorender.com.

## Author Contribution

J.B. and J.T.J. performed experiments with help from V.B., A.K.T., A.T., J.F. and W.N. J.Bo. performed AAV electroporations for TCR KI mouse line generations. S.D. and P.G.T. provided TCR sequences used for the TCR KI mice. F.S., H.H. and A.E. optimized and provided AAV plasmids. G.D.V., AM.B. and J.T.J. supervised the work. A.M.B. and J.T.J. conceptualized the work and designed experiments. J.B., A.B. and J.T.J. wrote the manuscript with input from all authors.

---

**Supplemental Figure 1 (related to Figure 2).**
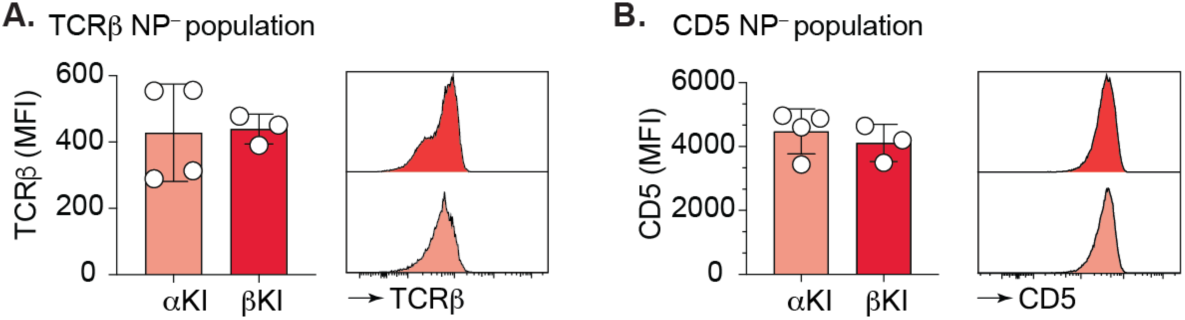
TCRβ and CD5 expression among escapees. Graphs and histograms show the expression of TCRβ (A) and CD5 (B) among NP-tetramer^−^ CD4^+^ thymocytes from the indicated TCR KI mice.

**Supplemental Figure 2 (related to Figure 3).**
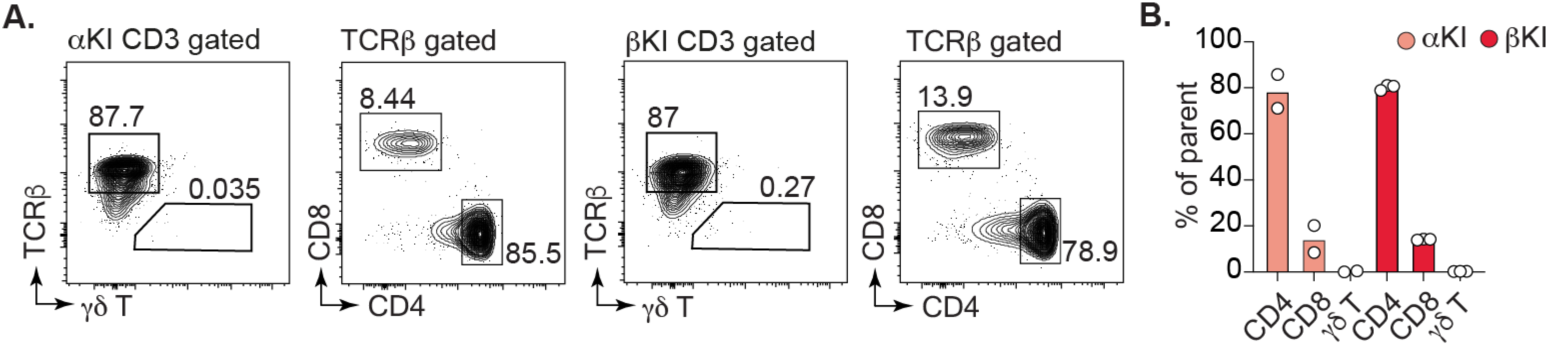
Frequency of γδ, CD4 and CD8 T cells in spleens of TCR KI mice. A) Representative flow cytometry plots indicating gating strategy for **γδ** CD8 and CD4 T cells. B) Quantification of major T cell subsets (**γδ** out of CD3^+^ T cells and CD4^+^ or CD8^+^ out of TCRβ^+^ T cells). Data show one experiment.

**Supplemental Figure 3.**
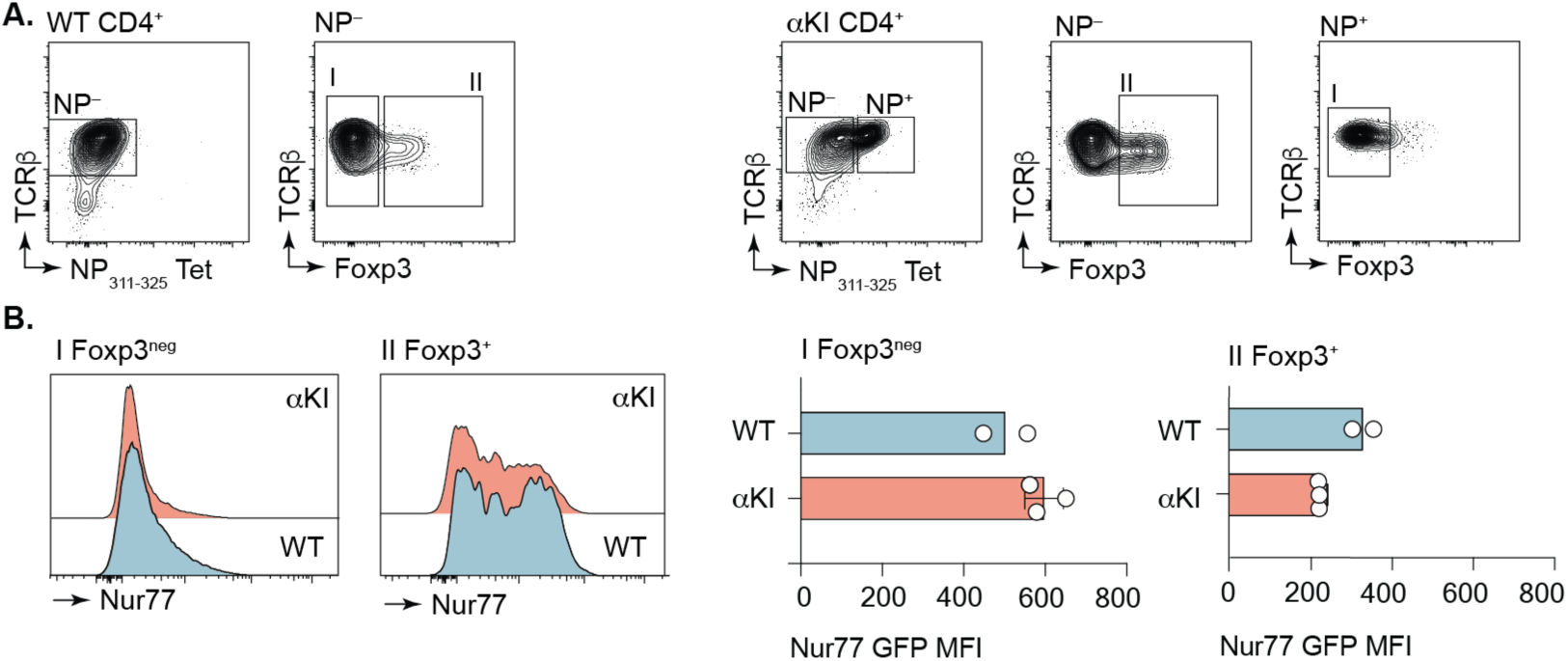
Nur77 expression in peripheral T cells. A) Representative plots showing Foxp3 expression in NP^+^ and NP^−^ populations in WT and αKI TCR mice crossed to Nur77-GFP reporter mice. B) Bar graphs show MFI for the Nur77 GFP fluorescence for NP-tetramer^+^ or NP-tetramer^−^ KI T cells and MFI of CD4^+^ WT T cells. Data show one experiment.

**Supplemental Figure 4.**
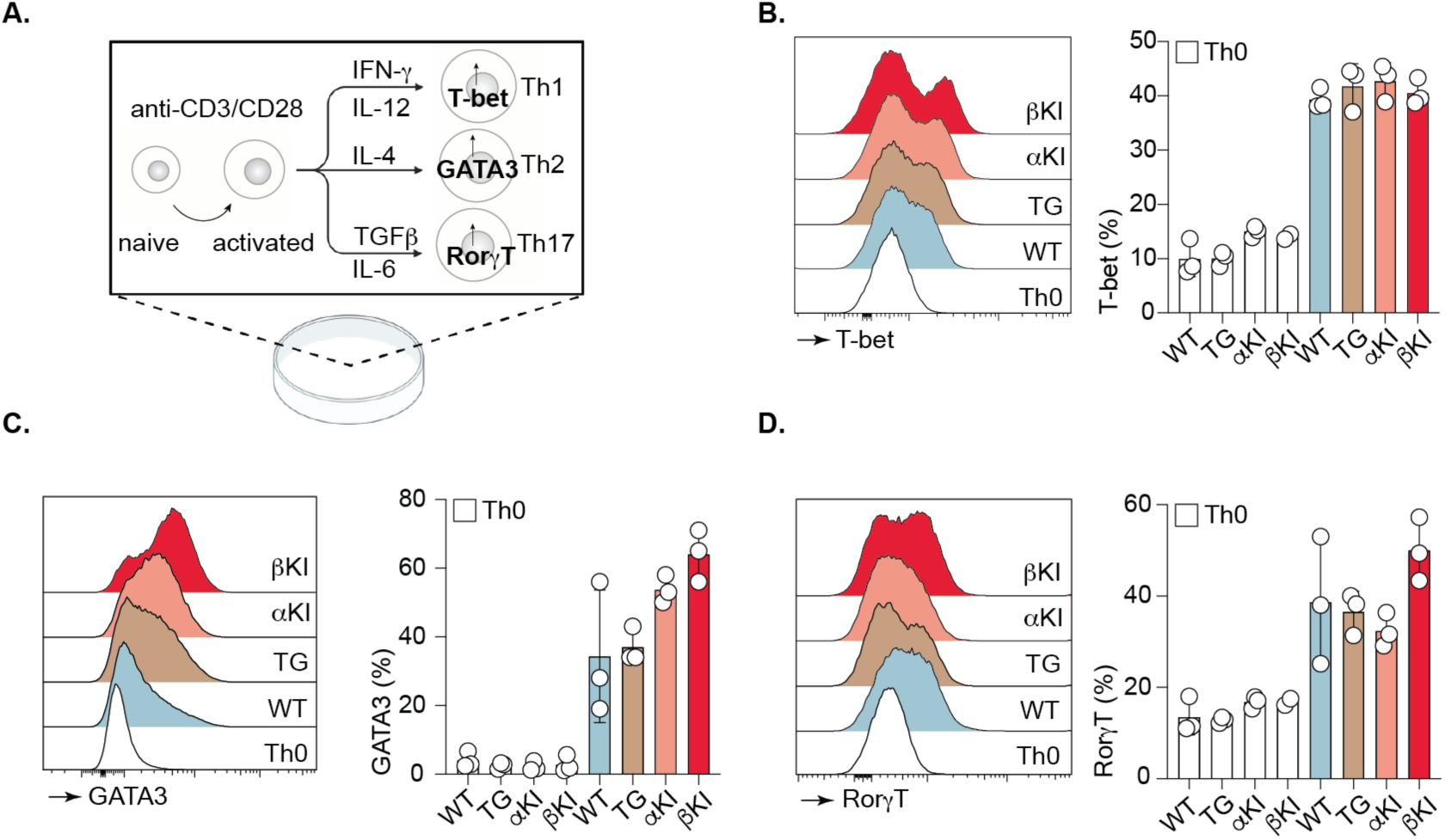
*In vitro* polarization of TCR KI T cells. A) Schematic showing *in vitro* polarization assay with stimulating and differentiating factors to obtain Th1, Th2 and Th17 with their respective transcription factors that were stained for and measured. Polarization of WT, TG, αKI and βKI T cells into Th1 B), Th2 C), and Th17 D) is summarized as mean ± s.e.m for T-bet, GATA3 and RorγT positive T cells respectively. Representative histograms are shown compared to Th0 for each condition. For all plots data and error bars represent the mean ± s.e.m. Data show one experiment.

**Supplemental Figure 5 (related to Figure 4).**
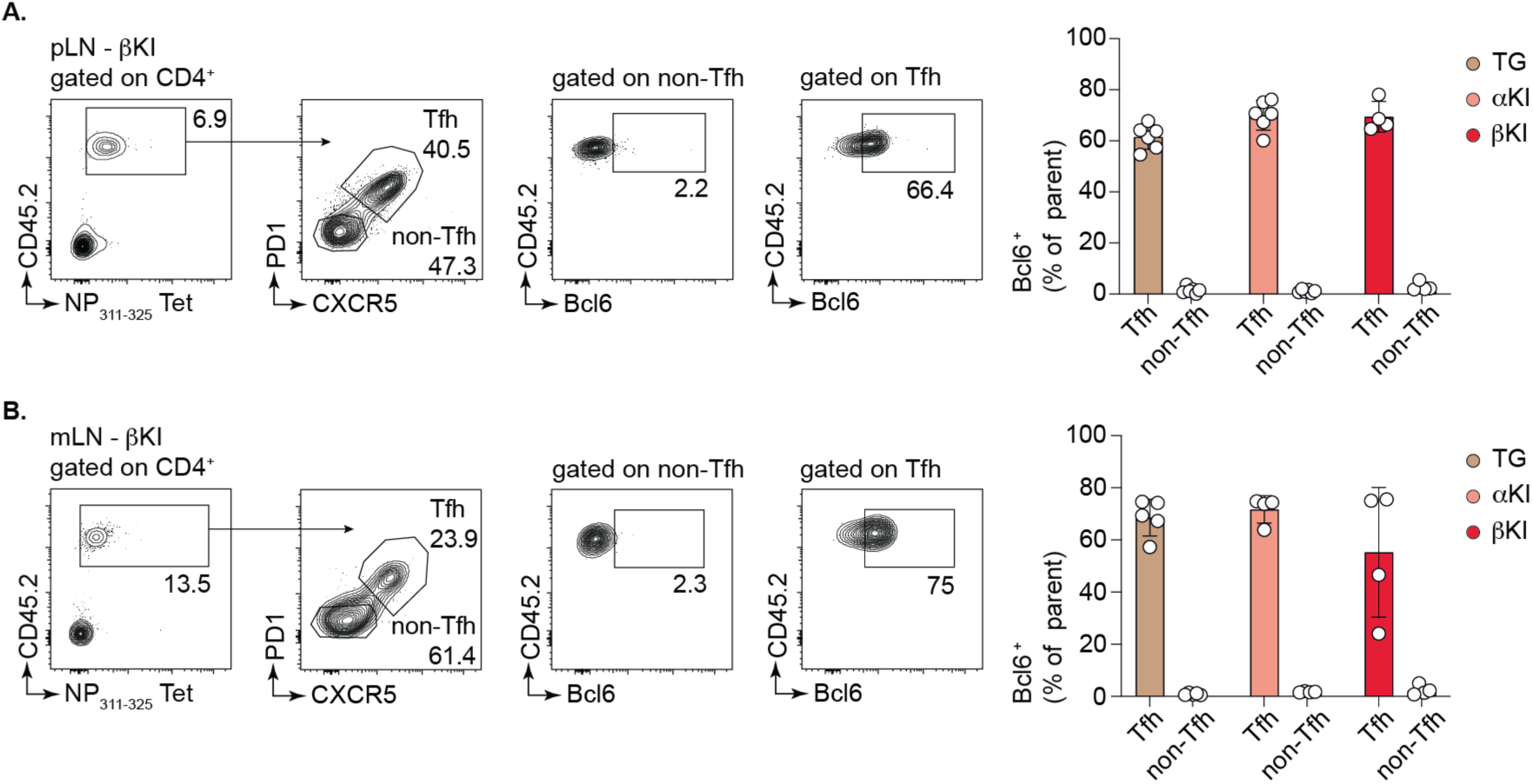
Bcl-6 expression in Tfh cells after protein immunization and PR8 infection. A) Representative flow cytometry plots from the pLN of a protein-immunized mouse showing gating strategy for Bcl6 expression gated on Tfh and non-Tfh of transferred CD4^+^ NP^+^ CD45.2^+^ T cells. Graph shows quantification. B) Representative flow cytometry plots from the mLN of a PR8-infected mouse showing gating strategy for Bcl6 expression gated on Tfh and non-Tfh of transferred CD4^+^ NP^+^ CD45.2^+^ T cells. Graph shows quantification. Data are pooled from two experiments. Data and error bars represent the mean ± s.e.m. pLN, popliteal lymph nodes; mLN, mediastinal lymph nodes.

